# Herpes Simplex Virus Infection, Acyclovir and IVIG Treatment All Independently Cause Gut Dysbiosis

**DOI:** 10.1101/844712

**Authors:** Chandran Ramakrishna, Stacee Mendonca, Paul M. Ruegger, Jane Hannah Kim, James Borneman, Edouard Cantin

## Abstract

Herpes simplex virus 1 (HSV) is a ubiquitous human virus resident in a majority of the global population as a latent infection. Acyclovir (ACV), is the standard of care drug used to treat primary and recurrent infections, supplemented in some patients with intravenous immunoglobulin (IVIG) treatment to suppress deleterious inflammatory responses. We found that HSV, ACV and IVIG can all independently disrupt the gut bacterial community in a sex biased manner when given to uninfected mice. Treatment of HSV infected mice with ACV or IVIG alone or together revealed complex interactions between these drugs and infection that caused pronounced sex biased dysbiosis. ACV reduced *Bacteroidetes* levels in male but not female mice, while levels of the Anti-inflammatory Clostridia (AIC) were reduced in female but not male mice, which is significant as these taxa are associated with protection against the development of GVHD in hematopoietic stem cell transplant (HSCT) patients. Gut barrier dysfunction is associated with GVHD in HSCT patients and ACV also decreased *Akkermansia muciniphila,* which is important for maintaining gut barrier functionality. Cumulatively, our data suggest that long-term prophylactic ACV treatment of HSCT patients may contribute to GVHD and potentially impact immune reconstitution. These data have important implications for other clinical settings, including HSV eye disease and genital infections, where ACV is given long-term.

**Author Summary:** Primary and reactivated HSV and VZV infections are treated with Acyclovir (ACV), an antiviral drug that blocks viral DNA synthesis. In some patients IVIG is used as adjunctive therapy to block deleterious inflammation. Long term preventative treatment of patients who receive stem transplants for various blood cancers has been successful in preventing life threatening reactivated HSV and VZV infections, but GVHD remains a major factor limiting transplant success. Studies reported here reveal that HSV infection, ACV and IVIG given alone can all disrupt the gut microbiota and that complex interactions between these drugs and infection results in even more pronounced sex biased changes in the gut bacteria community structure. Importantly, ACV treatment decreased the levels of specific bacterial taxa, including the anti-inflammatory *Clostriodia* and *Bacteroidetes* that have been shown to protect against development of GVHD in stem cell transplant patients. These data suggest that long term preventative treatment of patients with ACV may contribute to GVHD in transplant patients and have negative consequences in other HSV induced diseases treated long term with ACV. The health effects of long term ACV and IVIG treatments warrant further clinical studies.

## Introduction

Herpes Simplex Virus type 1 (HSV), a ubiquitous human virus is the major cause of HSV encephalitis (HSE), the most prevalent sporadic encephalitis resulting from either primary infection or reactivation of latent virus. However, despite improved diagnostic procedures and effective antiviral therapies, most HSE survivors have persistent neurological impairments, including memory and behavior disturbances, dysphasia and seizures, and only 50-65% of these survivors return to independent living [1, 2]. A delay in initiating Acyclovir (ACV) treatment past the second hospital day is associated with poor neurological outcomes [3, 4]. Recent clinical trials evaluating prolonged oral ACV/valaciclovir (VACV) treatment following standard 14-day intravenous ACV treatment reported improved neurocognitive outcomes in neonates but not adults for reasons that are obscure [5, 6]. Although, it is generally accepted that replication induced pathology underlies HSV related neurological dysfunction, supporting experimental or clinical evidence is lacking. Overwhelming evidence has linked inflammation to the development of various neurological disorders and neuropsychiatric diseases, including Alzheimer’s disease (AD), schizophrenia, autism spectrum disorder (ASD), multiple sclerosis (MS), Parkinson’s disease (PD), depression and anxiety [7–9].

Having unequivocally established that HSE arises from exaggerated CNS inflammatory responses and that the immunomodulatory activities of intravenous immunoglobulins (IVIG) can prevent HSE in a mouse model [10], we tested the hypothesis that persistent inflammation, which is documented in humans and mice after HSE [11–14], causes neurobehavioral impairments in survivors, that should be impeded by IVIG’s anti-inflammatory activity [10]. Compared to treatment of HSV infected mice with ACV or PBS alone, treatment with ACV+IVIG from day 4 pi reduced CNS inflammation and anxiety, consistent with our hypothesis. Strikingly, development of learning and memory (LM) deficits that were evident only in female PBS treated mice, were inhibited by ACV treatment and counterintuitively, aggravated by ACV+IVIG treatment. Treatment of infected male mice with ACV+IVIG also impaired LM compared to ACV or PBS alone, revealing that IVIG antagonized the beneficial effects of ACV [15]. Intriguingly, the differential antagonistic effects of ACV+IVIG on cognitive behavior in HSV infected mice, compared to ACV and PBS treatment alone, were reflected in differential serum proteomic profiles [15]. These reported antagonistic effects of ACV and IVIG on LM present a conundrum, since they are at odds with the known mechanisms of action of these drugs.

Rapidly accumulating evidence is revealing the critical role of the microbiome in regulating brain homeostasis and function such that perturbation of the gut bacteria community structure and function is increasingly being implicated in a variety of neurodegenerative and neuropsychiatric diseases. In an effort to gain insight into how HSV induces LM impairment and the paradoxical effects of ACV and IVIG, we investigated a role for the gut microbiota. HSV infection, ACV and IVIG were all associated with significant disruption of the gut bacterial community structure that was sex biased. Furthermore, treating HSV infected mice with either ACV or IVIG alone or both drugs together resulted in more pronounced sex-biased shifts in the gut bacterial community structure compared to uninfected mice. These results have significant clinical implications, particularly when patients receive prolonged ACV or IVIG treatment.

## Results

Equal numbers (n=8) of female and male C57BL/6 mice were bilaterally inoculated with virulent HSV1 strain 17+ (1×10^5^ PFU/eye) by corneal scarification as previously described [15]. At day 4 post infection (pi), ACV was administered at 1.25 mg / mouse by intraperitoneal injection (ip) daily for 3 days, while IVIG was given as single dose of 25 mg/mouse by ip injection on day 4pi [15]. Fresh fecal pellets (n=1-2/ mouse) were collected on day 7 pi and stored at −80°C until processed for Illumina 16S rRNA gene sequencing to determine the effects of infection and drug treatment on the gut microbiome. Normal male and female mice differed in gut bacteria composition and unexpectedly, HSV ocular infection caused further shifts in the gut bacteria community and amplified this sex difference, as shown in a PCoA plot of Hellinger beta diversity distance values for infected compared to uninfected male and female mice (**Figure 1A**; P<0.05, Adonis Tests). In addition, HSV infection had a greater effect on gut bacterial communities in males (P=0.003) compared to females (P=0.011) (**Figure 1A**). Significant differences were observed at the phyla level, particularly for firmicutes (**Figure 1B**) with more marked differences evident at the species level for *Clostridium aerotolerans* and other clostridial species, for example *Clostridium XIVa* that ferment carbohydrates in the gut resulting in production of short chain fatty acids (SFCs) that contribute to barrier integrity and also exhibit anti-inflammatory properties (**Figure 1C**). A notable difference was also observed for *Akkermansia muciniphila* that has many health promoting activities, including maintaining gut barrier health (**Figure 1C**).

**Figure 1.**
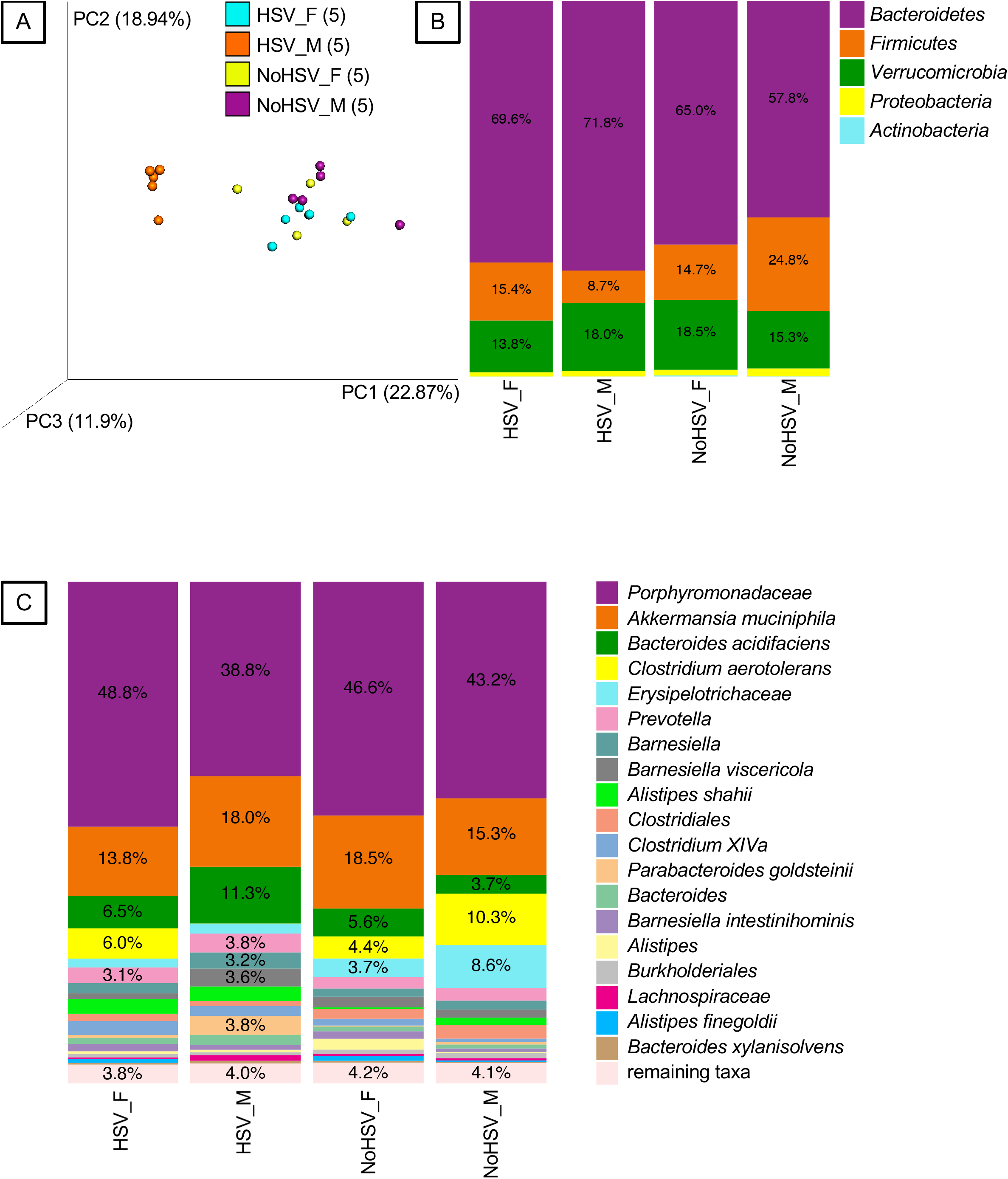
Fecal Bacteria from HSV-Infected and Uninfected Mice. A. Principal-coordinates analysis (PCoA) of Hellinger beta diversity distance values generated from 16S rRNA gene sequences. All four groups were different (P<0.05, Adonis Tests). The number of mice (n) in each genotype-microbiota group are shown in parentheses. B. Bacteria phyla associated with HSV-infected and uninfected mice. C. Bacterial species (or higher taxa) associated with HSV-infected and uninfected mice. Females = _F and Males = _M.

Treating HSV infected mice with ACV from day 4 pi for three days resulted in even more drastic shifts in the gut bacteria composition and exaggerated sex differences (**Figure 2A**), than for infection alone. Considerable abundance changes were evident at the Phyla level for *Bacteroidetes*, *Firmicutes* and *Verrucomicrobia* (**Figure 2B**) and at the species level (**Figure 2C**). Notably, whereas HSV infection reduced the abundance of *Firmicutes* significantly in male but not female mice (**Figure 1B**), ACV reversed this effect restoring the abundance to the level in uninfected male mice, while also increasing the abundance in female mice (**Figure 2B** and **Figure 1B**). Notable abundance changes at the species level included drastic suppression of *Clostridium aerotolerans* in infected male mice compared to increased abundance in females (**Figure 1C**), while ACV treatment further increased this abundance only in females (**Figure 2C**). *Akkermansia muciniphila* abundance was increased by infection in male mice but reduced in females (**Figure 1C**), while ACV treatment resulted in total suppression of this species in female mice compared to a marked reduction in male mice (**Figure 2C**). There are many other similar changes in species abundance that are differentially impacted by ACV treatment in a sex-biased manner, indicative of complex interactions between infection, ACV effects on infected host cells, and bacteria, as well as metabolites produced by bacterial metabolism of ACV.

**Figure 2.**
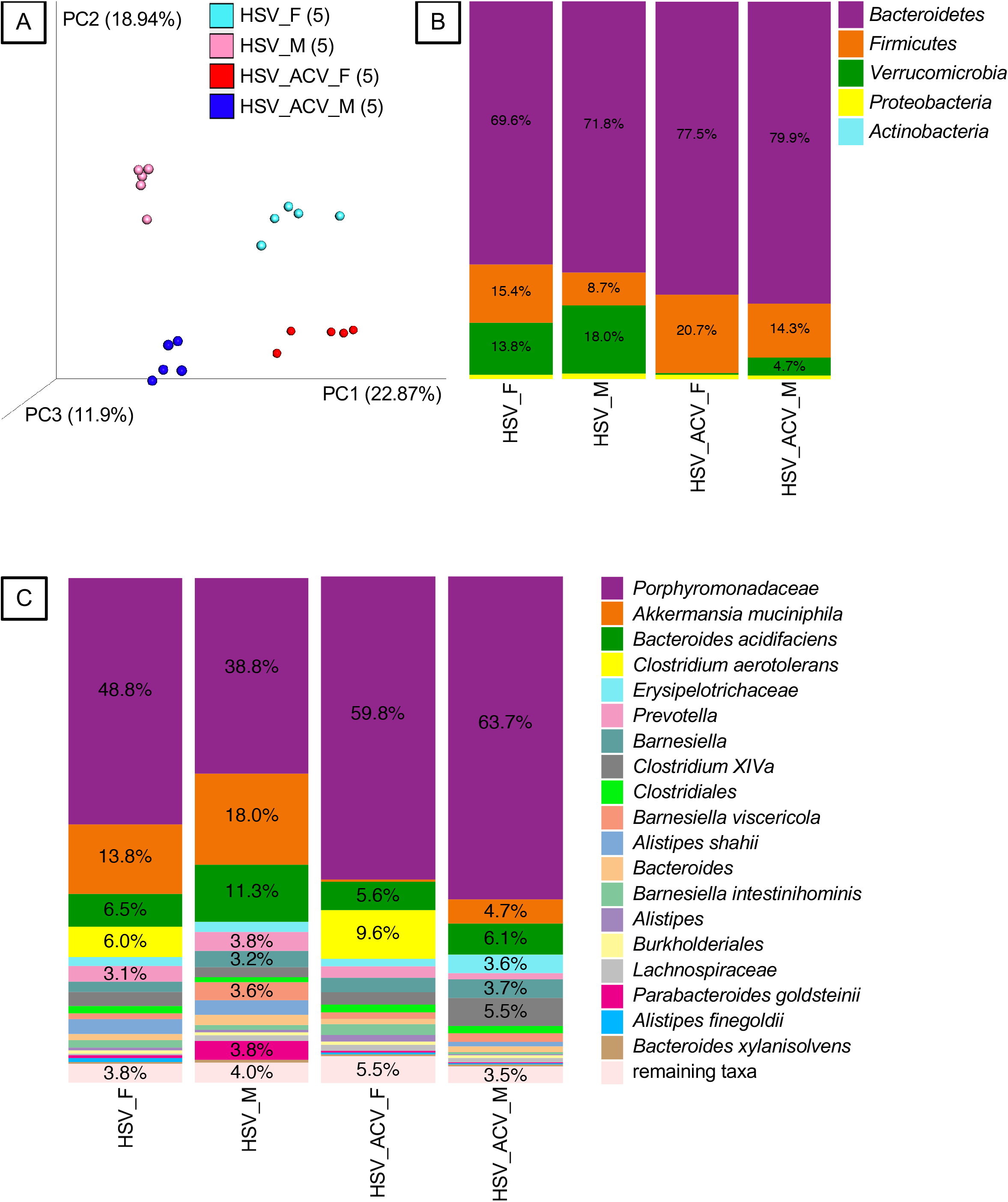
Fecal Bacteria from HSV-Infected Mice Treated and Not Treated with ACV. A. Principal-coordinates analysis (PCoA) of Hellinger beta diversity distance values generated from 16S rRNA gene sequences. All four groups were different (P<0.05, Adonis Tests). The number of mice (n) in each genotype-microbiota group are shown in parentheses. B and C. Bacteria phyla and species (or higher taxa), respectively, associated with HSV-infected mice treated and not treated with ACV. Females = _F and Males = _M.

Treatment of uninfected mice with IVIG alone also shifted the gut bacteria community composition with a notable marked sex effect as determined by a beta diversity analysis (**Figure 3**). Males and females showed a major reduction in *A. muciniphila,* and a lesser reduction of *Verrucomicrobia* in males, compared to females that showed increased abundance of this phylum in response to IVIG treatment (**Figure 4**). The abundance of many other bacterial species was differentially altered by IVIG treatment of males and females, for example, *Clostridium aerotolerans*, *Bacteroides acidifaciens* and *Porphyromonadaceae* (**Figure 4B**). The response to IVIG was distinct in HSV infected mice, and the complex interactions between infection, ACV and IVIG were also evident at the phyla and species levels and were strongly sex biased as well (**Figure 4A and 4B**). IVIG treatment decreased *A. muciniphila* abundance markedly in infected males and females as did ACV, whereas in contrast, treatment with ACV+IVIG caused a notable increase in its abundance, indicative of antagonistic effects of these two drugs in the context of infection (**Figure 4B**) In a similar vein, *C. aerotolerans* abundance increased markedly in males, but was unchanged in females treated with IVIG, while in contrast, it was strongly decreased in males but slightly increased in females treated with ACV alone. In contrast, treatment with ACV+IVIG suppressed an IVIG-induced increase in males and an ACV-induced increase in females, revealing antagonism between ACV and IVIG in the context of HSV infection (**Figure 4B**).

**Figure 3.**
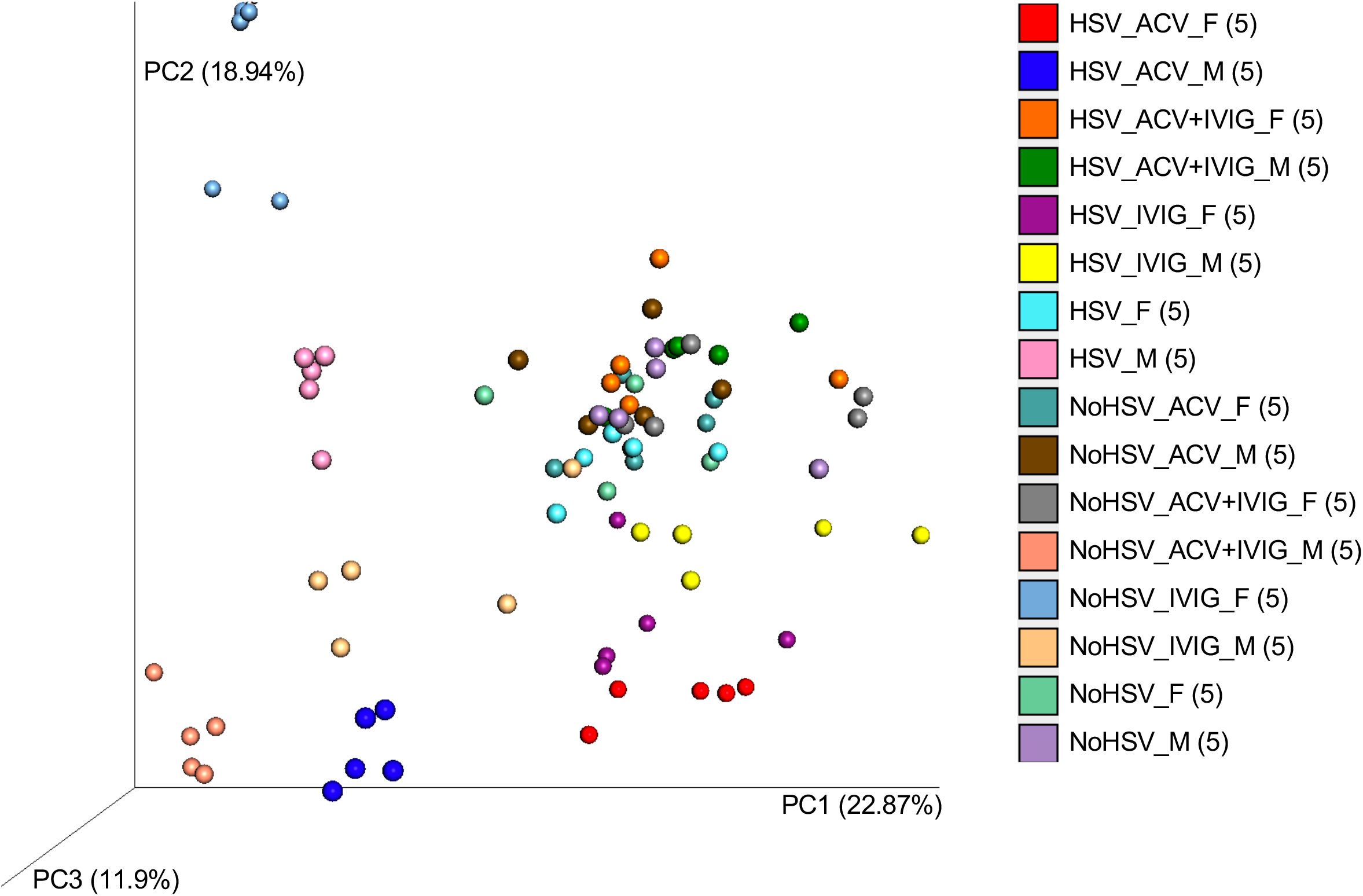
Beta Diversity Analysis of Fecal Bacteria from HSV-Infected and Uninfected Mice Treated and Not Treated with ACV and/or IVIG. Principal-coordinates analysis (PCoA) of Hellinger beta diversity distance values generated from 16S rRNA gene sequences. The number of mice (n) in each genotype-microbiota group are shown in parentheses. Females = _F and Males = _M.

**Figure 4.**
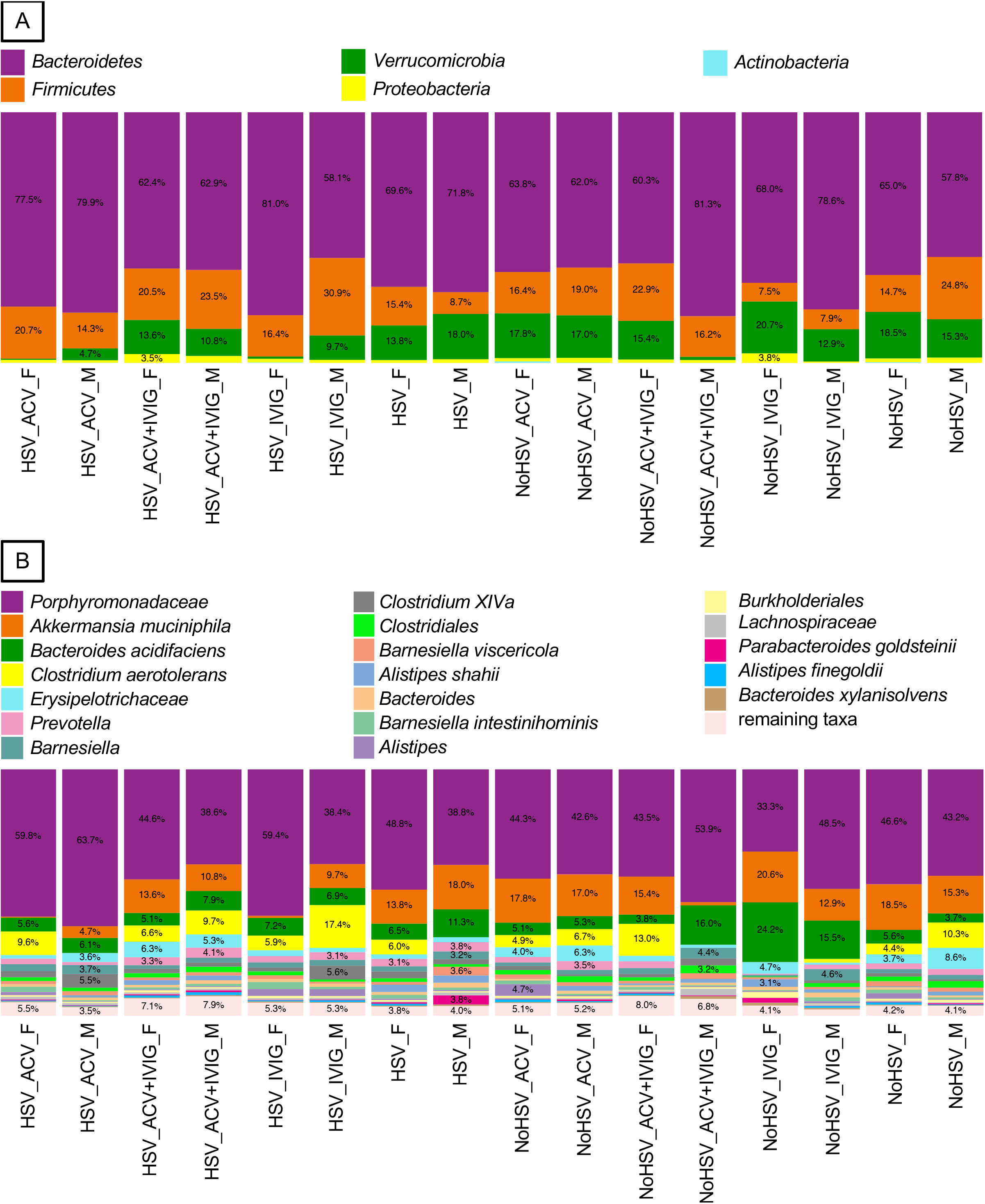
Fecal Bacterial from HSV-Infected and Uninfected Mice Treated and Not Treated with ACV and/or IVIG. A and B. Bacteria phyla and species (or higher taxa), respectively, associated with HSV-infected and uninfected mice treated and not treated with ACV, IVIG, or ACV+IVIG. Females = _F and Males = _M.

Patients with hematologic and other malignancies have benefited immensely from allogeneic hematopoietic stem cell transplantation (allo-HSCT or HSCT), which can be a potent curative immunotherapy. However, life threatening complications such as graft-versus-host disease (GVHD), relapse, and infections that include reactivated HSV and VZV limit its application [16]. HSV and varicella zoster (VZV) reactivation has been successfully suppressed by prophylactic ACV treatment, though ACV-resistant (ACVr) HSV is an emerging problem [17, 18]. Long term ACV prophylactic treatment is now routine for HSCT patients, because it was found to correlate with reduced HSV and ACVr HSV disease in those treated for longer than 1 year [19].

Given this routine clinical practice, we evaluated the effects of ACV on fecal bacteria, because gut microbes have been implicated in GVHD pathophysiology and because we posit that ACV contributes to the development of GVHD by changing the gut microbiota. First, we identified gut bacterial changes in humans with GVHD [20–30]. Next, we determined whether the ACV-induced changes that we detected in this mouse study matched those GVHD-associated changes. Whenever we identified taxa that were altered in both types of studies, the direction of the change was the same, and it was consistent with our hypothesis that ACV contributes to the development of human GVHD by changing the gut microbiota. In the following, we describe these results, and we note that these ACV-induced changes were only observed in the HSV-infected mice and not in the uninfected mice.

Reduced levels of several taxa belonging to the phylum *Bacteroidetes* have been shown to be associated with GVHD, indicating that these gut bacteria may play a protective role. In a pediatric study, GVHD patients had lower levels of the family *Bacteroidaceae* and the genus *Parabacteroides* [30]. In a longitudinal study, pediatric patients that had lower levels of *Bacteroidetes* prior to HSCT were more likely to develop GVHD [24]. In our study, all three if these taxa were reduced by ACV treatment in male but not female mice (**Figure 5A**).

**Figure 5.**
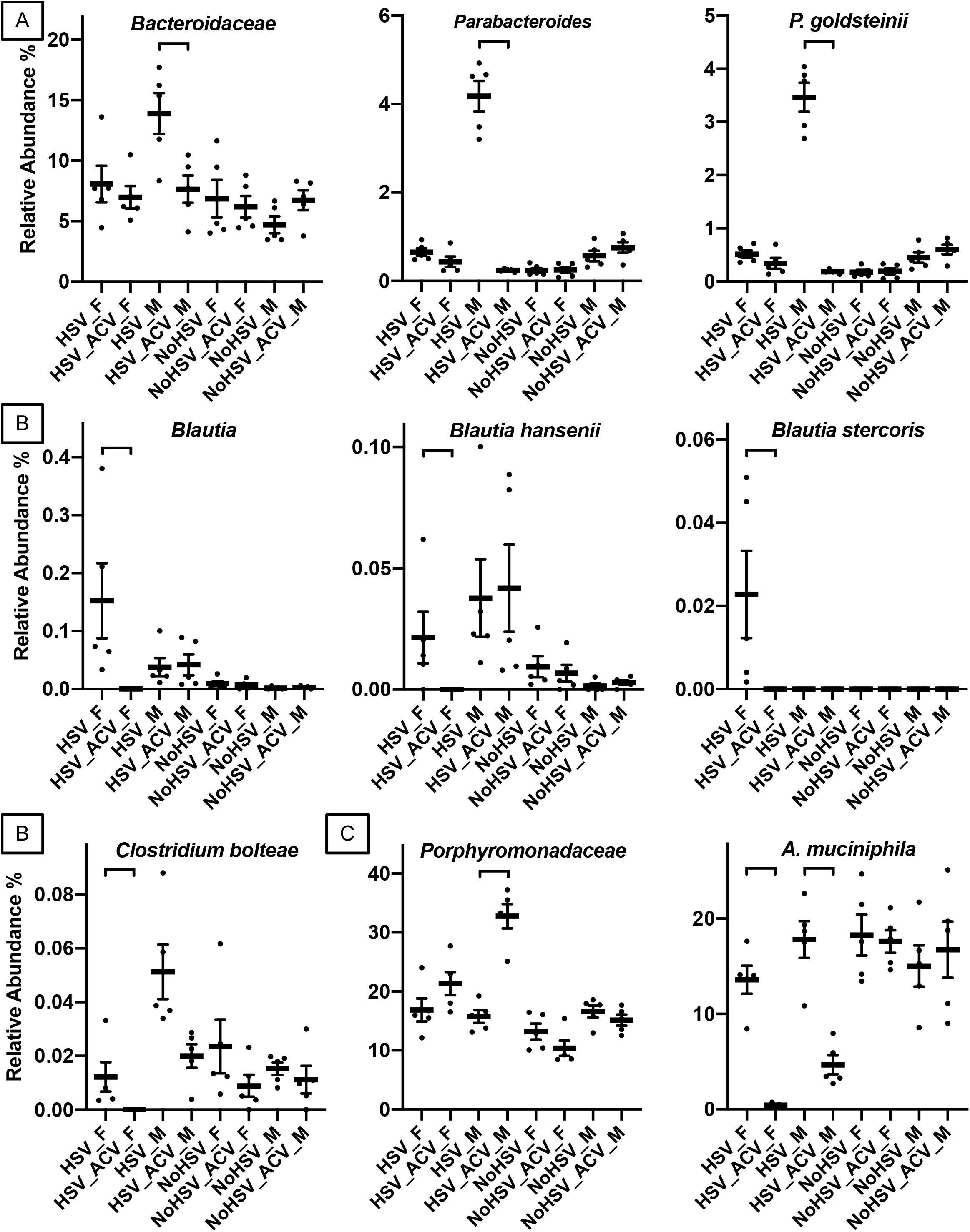
Fecal Bacterial from HSV-Infected and Uninfected Mice Treated and Not Treated with ACV. A and B. Fecal bacterial taxa that were were changed in both human GVHD studies and by ACV in this study. A and B. Members of the *Bacteroidetes* and AIC, respectively. C. The two most abundant bacterial OTUs. The only pairwise differences shown are between ACV treated and untreated mice for each sex (FDR-adjusted P values < 0.05). Bars = standard error. Females = _F and Males = _M.

Reduced levels of Anti-Inflammatory Clostridia (AIC) have also been detected in human GVHD patients [20, 23–25, 27–30], indicating that these gut bacteria may play a protective role. This terminology was first introduced by Piper et al. [31] in the context of short bowel syndrome, and then introduced to the GVHD literature by Simms-Waldrip et al. [30]. AIC taxa include members of the families *Clostridiaceae*, *Erysipelotrichaceae*, *Eubacteriaceae*, *Lachnospiraceae* and *Ruminococcaceae*. In a pediatric study, decreases in *Blautia* and *Clostridium bolteae* were associated with the development of GVHD [30]. In an adult study, lower levels of *Blautia*, *Blautia hansenii*, and *Blautia stercoris* were associated with the development of GVHD [28]. In a longitudinal study, reduced levels of the *Blautia* before HSCT was shown to be a predictive marker for the development of GVHD [27]. In our study, all of these taxa were reduced by ACV treatment in female but not male mice (**Figure 5B**).

In a more detailed analysis of AIC bacteria, we observed that while HSV infection increased the abundance of *Blautia hansenii* only in males, ACV treatment reduced its abundance in females but had no effect on its abundance in males (**Supplemental Figure 1**). Remarkably, a dramatic increase in *B. hansenii* in uninfected females was observed after IVIG treatment, and this increase was abrogated by ACV (compare NoHSV_F, NoHSV_IVIG_F and NoHSV_ACVplusIVIG_F) (**Supplemental Figure 1**), a result that supports sex-based differential effects of these drugs. However, during HSV infection, both IVIG and ACV reduced *B. hansenii* in females, whereas only IVIG reduced abundance in males. Interestingly, HSV infection significantly increased the abundance of the AIC genera *Blautia, Allobaculum,* and *Clostridium* XVIII but not *Turicibacter* in both males and females (**Supplemental Figure 2**). ACV treatment of HSV infected female mice resulted in significant decreases in the abundances of 4 AIC genera: *Blautia*, *Allobaculum*, *Clostridium* XVIII and *Turicibacter,* whereas in infected males, ACV decreased the abundance of *Marvinbryantia* and *Oscillibacter* (**Supplemental Figure 2**). In addition, ACV increased the abundance of *Turicibacter* in uninfected females but not males.

Finally, the two most abundant operational taxonomic units (OTUs), which exhibited a change in their relative abundances due to ACV treatment, were assigned to the family *Porphyromonadaceae* and the species *A. muciniphila* (**Figure 5C**). While we did not find these taxa associated with GVHD in prior human studies, GVHD has been associated with intestinal barrier dysfunction [32–36]. Supporting our hypothesis that ACV contributes to the development of GVHD by changing the gut microbiota, members of the *Porphyromonadaceae* have been shown to cause gut barrier dysfunction [37, 38], and our *Porphyromonadaceae* OTU was increased in its abundance by ACV. In addition, *A. muciniphila* was decreased by ACV treatment in our study, and it has been shown to strengthen gut barrier functioning [39–41].

## Discussion

Our intention in this brief report is to alert the scientific community and especially clinicians to the fact that HSV infection, the antiviral drug ACV, and the immunomodulatory biological, IVIG, can all independently result in significant perturbations of the gut bacterial communities. Our data reveal complex interactions between HSV infection and ACV or/and IVIG treatment that result in marked alterations to gut bacterial communities. Although the clinical consequences of these changes have not yet been elucidated, they could have profound implications in several settings including HSCT-associated GVHD.

Though the mechanisms by which ocular HSV infection causes gut dysbiosis are unclear, neuroinflammatory mechanisms and effects on the enteric nervous system via connected brainstem neuronal circuits can be envisaged [15, 42]. Indeed, recent paradigm-shifting reports reveal that peripheral neurons, including nociceptive and sensory neurons, can directly sense and respond to environmental alarms by releasing neuropeptides that can regulate immune responses in target organs including the gut [43, 44]. Persistence of gut dysbiosis was not evaluated here, but results from a behavioral study alluded to earlier suggest long-term effects of infection and drug treatment on gut bacterial ecology should be investigated [15]. Sex biased effects on HSV induced dysbiosis merit further study, as these may involve microglial responses to HSV infection and the microglial compartment is known to be regulated by the microbiota in a sex biased manner [45–47].

The mechanism by which ACV, the standard antiviral for HSV infections, changes the gut microbiota likely involves its uptake into bacteria. ACV is preferentially phosphorylated by the viral encoded thymidine kinase (Tk) resulting in cell retention and eventual incorporation into viral DNA resulting in inhibition of viral replication via DNA chain termination. Because Tk is conserved in numerous bacterial species, ACV can be taken up and incorporated into DNA, resulting in bactericidal effects [48–51]. Indeed, early studies on DNA replication mechanisms relied on labeling bacterial DNA with tritiated thymidine and many bacterial taxa can be imaged using nucleoside analogues such as 1-(2_-deoxy-2_-fluoro-_-D-arabinofuranosyl)-5-[125I] iodouracil ([125I]FIAU) that are substrates for HSV Tk [52–55]. Incorporation of [*methyl*-^3^H]thymidine into DNA has been unequivocally demonstrated for members of the *Clostridium* genus [56] and our data show ACV reduced the abundance of the *Blautia* genus (order *Clostridiales;* [57]) *Blautia hansenii*, *Blautia stercoris*, and *Clostridium bolteae* in females but not males. Additionally, interrogating the NCBI reference genome sequence for *Blautia hansenii* confirmed the presence of a thymidine kinase enzyme. Our data are therefore consistent with ACV causing dysbiosis by, at least in part, inhibiting the growth of various bacteria taxa via the Tk mechanism, though other mechanisms involving bacterial metabolism of ACV cannot be excluded. Clearly, the mechanisms by which ACV affects gut bacterial ecology are complex, which is further supported by the sex-biased effects.

We also explored the effects of IVIG treatment alone and in combination with ACV in HSV-infected and uninfected mice, because IVIG has been used to treat HSV encephalitis (HSE) and is also a frontline therapy for autoimmune encephalitis, which is triggered by HSE and other insults [58–60]. Moreover, IVIG is being evaluated in a randomized control trial for children with all-cause encephalitis to determine whether neurological outcomes are improved compared to standard antiviral therapy alone, which is similar to our behavioral study that generated paradoxical results [15, 61]. Reports that IVIG’s antigenic repertoire includes reactivities to a variety of gut commensal antigens and metabolites have increased recently [62–64], which is consistent with a report that gut commensals can somehow trigger systemic IgG responses under homeostatic conditions that protect against systemic infection [65, 66]. We speculate that by neutralizing bacterial/host antigens/metabolites, IVIG is able to influence host immunity, the nervous system, and other physiological processes, resulting in perturbation of gut bacteria ecology. We speculate that the disparate and complex effects of ACV and IVIG alone and in combination on the gut bacteria ecology likely account for their antagonistic effects on cognitive behavior in mice latently infected with HSV that we alluded to earlier [15].

This study has several limitations. Being exploratory in nature, analyses of the gut bacteria were done at a single time point immediately after infection or drug treatment, rather than as a longitudinal study that would have provided information on the persistence of the dysbiotic state as well as mechanistic insights as to how HSV, ACV and IVIG provoke dysbiosis. Ideally, the effects of ACV should be tested in latently infected mice, since virtually all HSCT patients harbor latent HSV. However, because HSV infection alone disrupts the gut bacterial community, assessing the effects of ACV on the gut bacteria community structure in the latently infected mice would likely be difficult. Because ACV was given ip to mice but usually orally to HSCT patients [67], its effects on the gut bacteria community maybe underestimated in our study.

Notwithstanding these caveats, our finding that ACV treatment of HSV infected mice decreased the relative abundances of several bacterial taxa is important because these bacteria have been negatively correlated with the induction of and mortality from GVHD in HSCT patients [24, 27, 28, 30]. These results are also consistent with our hypothesis that ACV contributes to the development of GVHD by changing the gut microbiota. In the context of allo-HSCT, GVHD occurs when donor immune cells recognize recipient tissues as foreign, leading to immune-mediated damage to several organs and tissues including the gastrointestinal tract. This has led researchers to posit that the reduction of anti-inflammatory bacteria such as AIC contribute to GVHD pathology [30]. The results from our study extend this hypothesis to include ACV treatment as a putative contributor to GVHD, because ACV reduced AIC bacteria in the gut. ACV treatment also decreased the relative abundances of several members of the *Bacteroidetes*, some of which have been shown to exhibit anti-inflammatory properties [68–71]. More relevantly, the capsular polysaccharide A (PSA) from *Bacteroides fragilis* reduced HSV-associated mortality in mice by dramatically reducing immune-mediated inflammation [72]. In addition, the two most abundant OTUs identified in our study, whose relative abundances were positively (*Porphyromonadaceae*) and negatively (*A. muciniphila*) correlated with ACV treatment, have been shown to weaken [37, 38] and strengthen [39–41] gut barrier function, respectively. These results provide an additional link between ACV treatment and GVHD, because barrier dysfunction, which can cause systemic inflammation, is a hallmark of GVHD [32–36]. Finally, long-term ACV prophylaxis initiated early after HSCT might also impair immune reconstitution based on results from a study of antibiotic depletion of gut bacteria in a murine model of syngeneic bone marrow transplantation [73]. These tantalizing results warrant independent validation and further detailed studies using a murine autologous BMT model to more rigorously evaluate the impact of long-term ACV prophylaxis on GVHD and engraftment, because results from such studies might eventually lead to improved outcomes for HSCT patients. Ideally, such future studies should be performed with mice harboring wild microbiota, because several recent reports show that immune responses in mice with wild microbiomes model human immune responses more closely than conventional mice with SPF microbiota [74–76].

## Materials and Methods

### Ethics Statement

All animal procedures were performed with prior approval of the City of Hope Institutional Animal Care and Use Committee (IACUC) under protocol # 07043 and within the framework of the Guide for the Care and Use of Laboratory Animals. C57BL6/J (B6) were bred in the vivarium at City of Hope.

### Mouse Studies

Master stocks of HSV1 strain 17 composed of only of cell-released virus were prepared in and their titers determined on mycoplasma-free CV-1 cell monolayers. Single use aliquots of virus in Hanks balanced salt solution supplemented with 2% fetal bovine serum were stored at −80°C. Male and female mice, 6–8 weeks of age, were infected with HSV1 17^+^, a virulent strain. Mice were sedated with ketamine (60 mg/kg) and xylazine (5 mg/kg) prior to HSV inoculation by corneal scarification. B6 mice were bilaterally inoculated with 1× 10^5^ PFU per eye and monitored daily as previously described [15, 77].

### Administration of Acyclovir and Intravenous Immunoglobulins

ACV obtained from (APP Pharmaceuticals, Schaumburg, IL) was given at 50 mg/kg of body weight by intraperitoneal (ip) injection daily for 3 days starting on day 4 pi and PBS was given according to the same schedule to control mice. IVIG (Carimune, NF) obtained from CSL Behring (King of Prussia, PA, USA) was given ip as a single 0.5 ml dose (25 mg/mouse) on day 4 pi or it was given in combination with a 3 day course of ACV.

### Illumina Bacterial 16S rRNA gene sequencing

Illumina bacterial 16S rRNA gene libraries were constructed as follows. PCRs were performed in an MJ Research PTC-200 thermal cycler (Bio-Rad Inc., Hercules, CA, USA) as 25 μl reactions containing: 50 mM Tris (pH 8.3), 500 μg/ml bovine serum albumin (BSA), 2.5 mM MgCl_2_, 250 μM of each deoxynucleotide triphosphate (dNTP), 400 nM of the forward PCR primer, 200 nM of each reverse PCR primer, 1 μl of DNA template, and 0.25 units JumpStart Taq DNA polymerase (Sigma-Aldrich, St. Louis, MO, USA). PCR primers 515F (GTGCCAGCMGCCGCGGTAA) and 806R (GGACTACHVGGGTWTCTAAT) were used to targeted the 16S rRNA gene containing portions of the hypervariable regions V4 and V5, with the reverse primers including a 12-bp barcode [78]. Thermal cycling parameters were 94°C for 5 min; cycles of 94°C for 20 s, 50°C for 20 s, and 72°C for 30 s, and followed by 72°C for 5 min. PCR products were purified using the MinElute 96 UF PCR Purification Kit (Qiagen, Valencia, CA, USA).

### 16S rRNA gene data processing

We used the UPARSE pipeline for de-multiplexing, length trimming, quality filtering and operational taxonomic units (OTU) picking using default parameters or recommended guidelines that were initially described in [79] and which have been updated at https://www.drive5.com/usearch/manual/uparse_pipeline.html. Briefly, after demultiplexing, sequences were trimmed to a uniform length of 249 bp, then filtered at the recommended 1.0 expected error threshold. Sequences were then dereplicated and clustered into zero-radius OTUs using the UNOISE3 algorithm [80], which also detects and removes chimeric sequences; this method is based on making OTUs at 100% identity. An OTU table was then generated using the otutab command. OTUs having non-bacterial DNA were identified by performing a local BLAST search [81] of their seed sequences against the nt database. OTUs were removed if any of their highest-scoring BLAST hits contained taxonomic IDs within Rodentia, Viridiplantae, Fungi, or PhiX. Taxonomic assignments to the OTUs were performed with SINTAX [82] using RDP Classifier 16S training set number 16 [83] as the reference database.

### 16S rRNA gene data analyses

Beta diversity was measured using QIIME 1.9.1 [84] to calculate a Hellinger beta diversity distance matrix, which was depicted using principle coordinates analysis (PCoA), and statistically assessed by performing Adonis tests. Statistical differences among the taxa were determined using edgeR [85, 86]. Taxa relative abundance figures were made using Prism (GraphPad, La Jolla, CA). Comparative analyses of the bacterial taxa between human GVHD studies and our mouse study excluded sequence-selective qPCR, because the selectivity of such assays is questionable given the conserved nature of the 16S rRNA gene, and because the results of such studies are not typically validated by sequence analyses. The bacterial sequences have been deposited in the National Center for Biotechnology Information (NCBI)’s Sequence Read Archive (SRA) under the BioProject Accession Number PRJNA549765.

