## Supplementary Fig. 1 and 2 for "Herpes Simplex Virus Infection, Acyclovir and IVIG Treatment All Independently Cause Gut Dysbiosis"

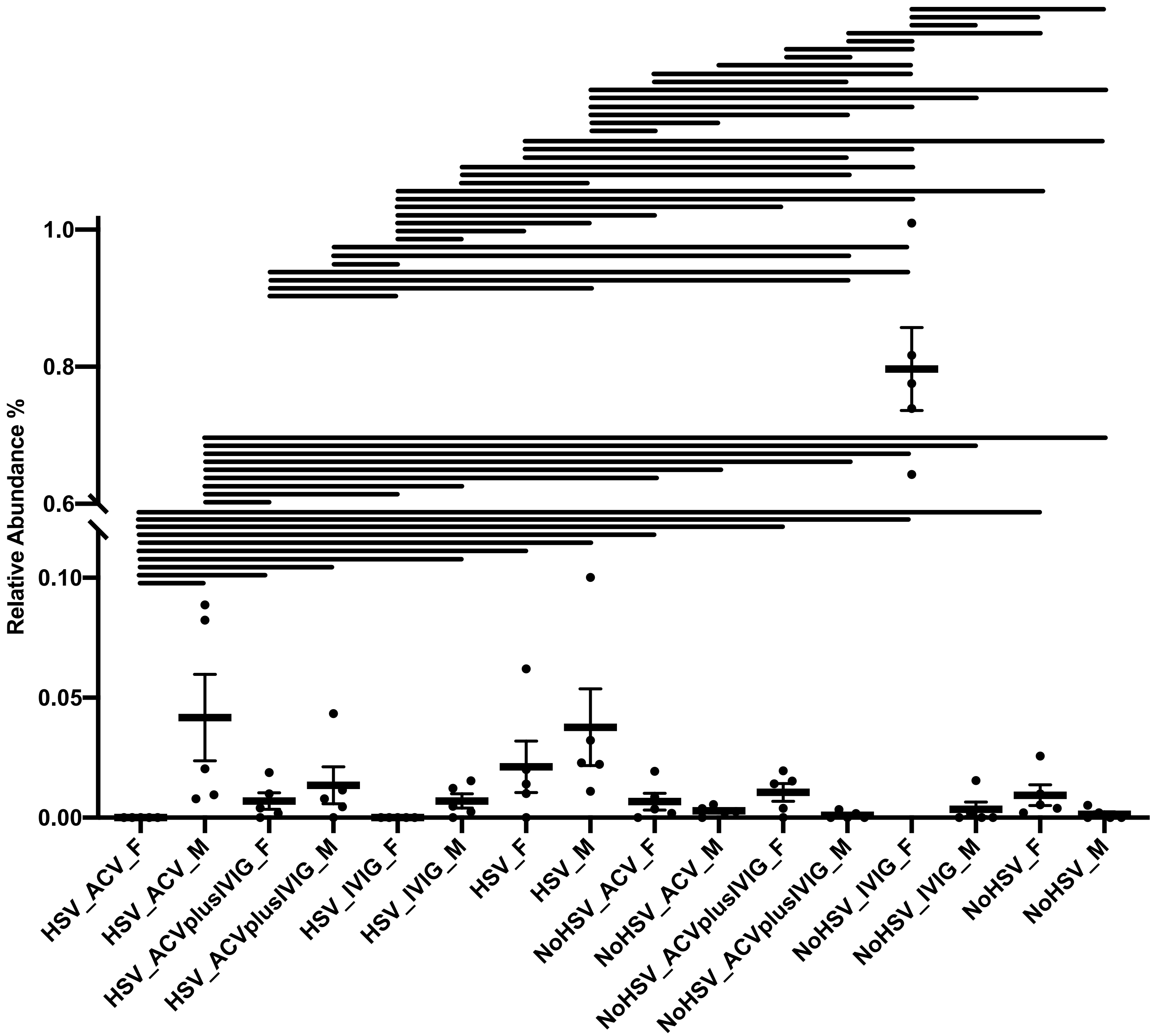


**Supplementary Figure 1. Fecal *Blautia hansenii* from HSV-Infected and Uninfected Mice Treated and Not Treated with ACV and/or IVIG**. Pairwise differences are shown by horizontal lines (FDR-adjusted P values < 0.05). Bars = standard error. Females = _F and Males = _M.

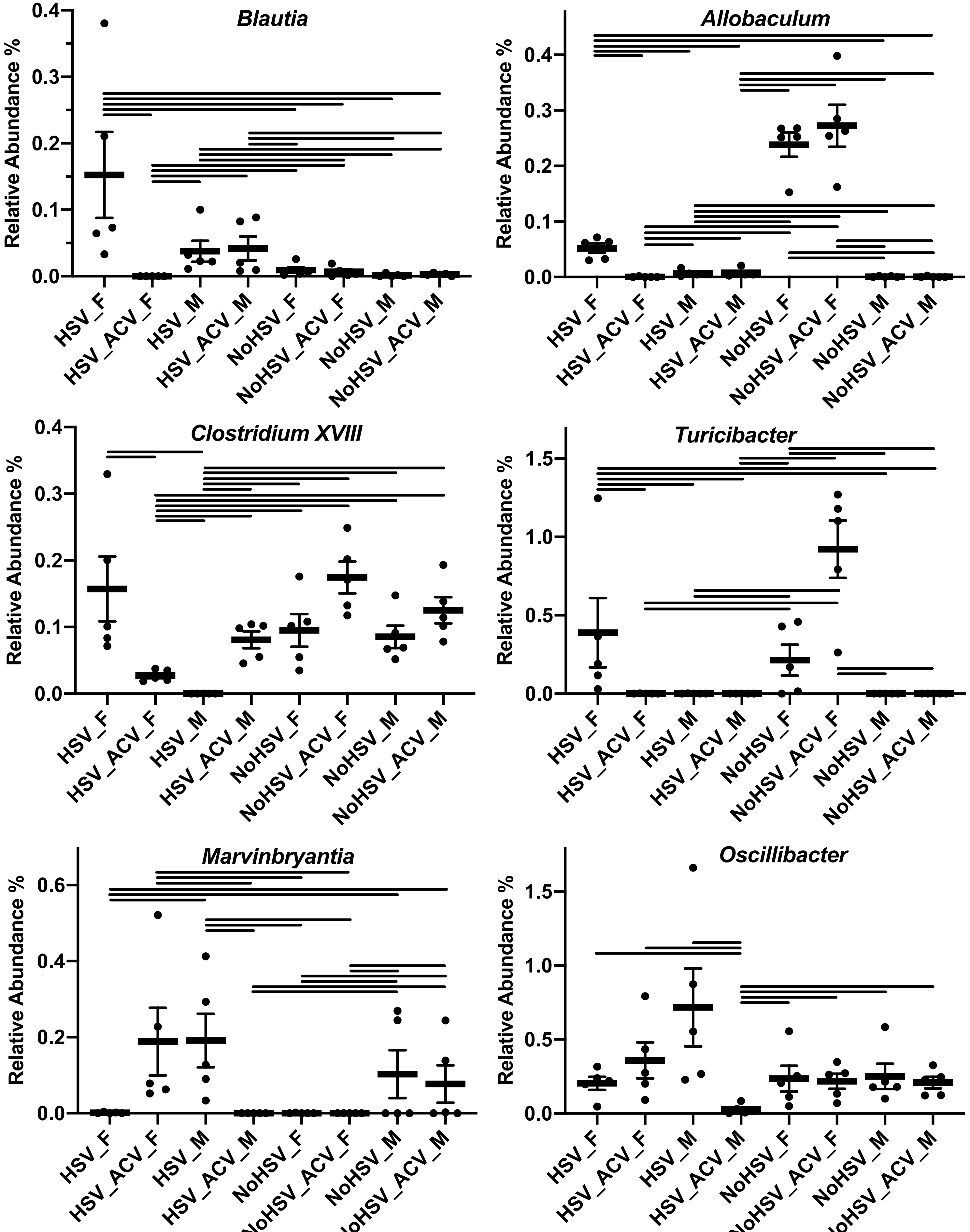


**Supplementary Figure 2. Fecal Bacterial Genera from HSV-Infected and Uninfected Mice Treated and Not Treated with ACV and/or IVIG**. Pairwise differences are shown by horizontal lines (FDR-adjusted P values < 0.05). Bars = standard error. Females = _F and Males = _M.
